# CombiCTx: Screening diffusion gradients of anti-cancer drug combinations

**DOI:** 10.1101/2025.02.27.635465

**Authors:** Christina Stelzl, Ada Lerma-Clavero, Selina Camenisch, Benoit Simon, Olle Eriksson, Oliver Degerstedt, Hans Lennernäs, Femke Heindryckx, Johan Kreuger, Paul O’Callaghan

## Abstract

The reduced effectiveness of chemotherapy in many patients highlights the need for novel drug combinations and optimal ratios that target multiple survival mechanisms, which tumors may engage to confer drug resistance. Dynamic conditions within the tumor microenvironment shape cell behavior and influence the response to anti-cancer drugs, varying by cell type and local context. Accordingly, assays that identify effective concentrations and drug interactions (additive, synergistic, or antagonistic) in a relevant tumor tissue model are required to discover new combination treatments. To address this need for combinatorial chemotherapeutic (CTx) screening assays, we reconfigured CombiANT, a device for testing antibiotic interactions, and present a new assay called CombiCTx. The assay uses a device with three reservoirs containing gels loaded with chemotherapeutics or other anti-cancer drugs. The drug-loaded device is inverted and placed in a standard culture dish containing cancer cells, and both are then enclosed in gel. As drugs diffuse from the reservoirs, cancer cells are exposed to overlapping dynamic gradients of anti-cancer drugs, which can interact in various ways. We imaged doxorubicin diffusion in the assay using timelapse microscopy and employed the apoptotic agent staurosporine as a model drug, and established an imaging protocol for quantifying MDA-MB-231 breast cancer cell apoptosis along drug diffusion gradients. Finally, we tested navitoclax and gemcitabine to demonstrate the capacity of CombiCTx to evaluate combined cytotoxic effects while accounting for drug diffusion. Evaluating drug combinations with an assay that accounts for drug properties that influence diffusion in a complex tumor matrix may provide clinically relevant information.

## Introduction

The last decades have seen a surge in the development of anti-cancer drugs targeting specific molecules and pathways required for cancer cell survival and cell division (1, 2). However, tumor heterogeneity can impair the full potential of such targeted therapies as the inherent instability of cancer cell genomes permits *de novo* mutations that compensate for the loss of the therapeutic target and promote drug resistance (3). It is therefore necessary to identify novel combination therapies, where two or more drugs that act by different mechanisms or modes of action are combined in an optimal dose ratio to reduce the probability that resistance will develop quickly (4). However, the development of combination therapies remains a major challenge, as treatment possibilities and synergism may be limited for example by treatment-induced toxicity, receptor antagonism and/or drug-drug interactions (4). It has also proven difficult and time-demanding to predict potent drug combinations (5); therefore, developing assays specifically designed to evaluate drug interactions can assist in identifying such beneficial combinations, and generate datasets that may prove useful in ongoing machine learning efforts to improve the accuracy of *in silico* predictions (6, 7). In addition to the benefits of combining treatments for additive effects, it is also valuable to identify potential synergistic and antagonistic effects of different treatment combinations (8). Effective drug synergisms permit the use of lower individual drug doses and thereby reduce the risk of side-effects, but it is well-known that the translation to patients is challenging (8–10). However, while a recent study evaluating the potency and efficacy of 2,025 two-drug combinations of clinical interest on 125 breast, colorectal and pancreatic cancer cell lines, demonstrated the benefits of combining different therapies, synergies were only detected in 5.2% of the tested cancer cell lines with drug combination pairs (11). Treatment of heterogenous tumors with combinations of effective drugs can ensure that subpopulations insensitive to one drug will be targeted by the other, and for many clinically approved combinations efficacy benefits are predominantly attributed to additive rather than synergistic effects (9).

Cellular responses to drug treatments, are context-dependent and modulated by a host of factors such as the composition and mechanical properties of the surrounding extracellular matrix (ECM), the concentration of solutes and gases, and pH, as well as by interactions with adjacent cells and the presence of different cell types in the tissue microenvironment (6, 12, 13). Several assays used for *in vitro* drug screening to identify possible combination therapies are conducted using microwell plates where cells are cultured on traditional two-dimensional plastic surfaces, and the efficiency of such systems comes at the expense of not being able to account for the complexity of the tumor microenvironment. However, there are ongoing efforts to develop scalable 3D assay systems that better recapitulate aspects of the tumor microenvironment with the aim of more efficiently identifying promising combination therapies (6, 14–19). While the potential patient benefits of drug combinations that have established synergies can be evaluated in clinical trials, the initial screening and identification of such synergies is heavily dependent on validated *in vitro* assays or well-established animal models (20).

In the current study we have developed CombiCTx, an anti-cancer drug screening assay. Cancer cells are encased in a hydrogel into which the CombiCTx device is embedded. The device consists of three reservoirs preloaded with anti-cancer drugs prepared in the same hydrogel, from which the drugs diffuse and create dynamic gradient landscapes over the entire cell population. We describe a detailed protocol on how to conduct the CombiCTx assay, and demonstrate an application to quantify diffusion of doxorubicin (DOX). We establish a time-lapse fluorescence imaging protocol to record apoptosis and cell death of MDA-MB-231 breast cancer cells, using a caspase 3/7 reporter and propidium iodide, and quantify responses to the apoptotic agent staurosporine in specific regions of the CombiCTx drug diffusion landscape. Finally, we apply this protocol and use CombiCTx to conduct a drug combination study using the anti-cancer agents navitoclax and gemcitabine.

## Materials and Methods

### Design and fabrication of the CombiCTx insert

The CombiCTx insert was designed using Autodesk Fusion 360 (Autodesk, San Francisco, USA) computer-aided design (CAD) software. Prototype CombiCTx inserts were 3D-printed using a Prusa i3 MK3S+ printer (Prusa Research, Prague, Czech Republic) with white X-PLA 1.75 mm filament (Add North 3D AB, Ölsremma, Sweden), which facilitated the testing of various design iterations. Prior to use the inserts were sterilized by extensive spraying with 70% ethanol, followed by 30 min of UV irradiation in a cell culture hood. These printed inserts were designed to fit into a 35 mm diameter cell culture dish and each reservoir holds a volume of 100 µl. The finalized CombiCTx insert design was milled from 8 mm thick sheets of polycarbonate using a CNC machine (Datron neo, Datron, CA, USA). This approach permits the production of a large numbers of inserts, and requires no post-processing. Prior to use these inserts were sterilized by autoclaving, and while relatively similar in dimensions to the 3D-printed inserts, had a larger reservoir volume of 124 µl.

### Cell culture

The MDA-MB 231 breast cancer cell line was obtained from the American Type Culture Collection (ATCC, LGC Standards GmbH, Wesel, Germany). The genetic identity was validated (21) by Idexx Bioanalytics (Ludwigsburg, Germany). Cells were grown in Dulbecco’s modified Eagle’s medium (DMEM) Glutamax (Thermo Fisher Scientific, Uppsala, Sweden), supplemented with 10% fetal bovine serum (FBS) (Thermo Fisher Scientific), and routinely grown and passaged before reaching confluency, at 37°C and 5% CO_2_ in a humidified incubator. For the various drug assays, cells were seeded to a density of 55 000 cells/cm^2^ in three wells of a tissue culture treated plastic 6-well plate one day before the start of the assay to ensure a near confluent cell layer.

### The CombiCTx assay setup

The CombiCTx insert consists of three reservoirs, which can be loaded with drugs of interest that are diluted in a low melting point agarose (LMPA, Thermo Fisher Scientific, Uppsala, Sweden) solution prepared with phenol red-free OptiMEM reduced serum medium (Thermo Fisher Scientific, Uppsala, Sweden). The insert is ready to be used once the LMPA has polymerized in the reservoirs. To activate a CombiCTx assay cell media is removed from a culture dish (35 mm in diameter) in which the cancer cells of interest had been seeded and cultured in advance. The cells are covered with LMPA (700 μl; 37°C), and thereafter the inverted CombiCTx is immediately inserted into the culture dish to activate the assay. As the cell-covering LMPA polymerizes it embeds the drug-containing LMPA reservoirs of the CombiCTx insert forming a continuous gel through which diffusion can occur (Fig. 1). The inverted insert is lowered into the culture dish at an angle such that two of the support pillars first touch the plate, followed by gently lowering the remaining pillar of the insert into place. This ensures minimal disturbance of the cell layer and avoids the trapping of air beneath the insert. To ensure the inserts remain stable during the polymerization process, a weight (stainless steel disc) is placed on top of the insert during polymerization, which typically occurred within 10-15 min. Media (1 ml) is then added to the culture dish to ensure all exposed surfaces are submerged, whereafter the assay is moved to the microscope for imaging, which is typically initiated within 10 min of activating the assay. We identified the LMPA-embedding of the loaded CombiCTx device in the cell culture dish to be a critical step. The objective of this process is to establish a continuum between the pre-polymerized LMPA in the reservoirs with the LMPA in the culture dish. Therefore, it is important not to disturb the setup during the polymerization process, and to subsequently handle the assay cautiously so as not to disturb the device in the gel, which could interrupt the interface between the hydrogels or introduce tears. Such imperfections may introduce channeling effects that will impact the diffusion from the reservoirs and we attributed a number of assay failures to such issues.

**Figure 1.**
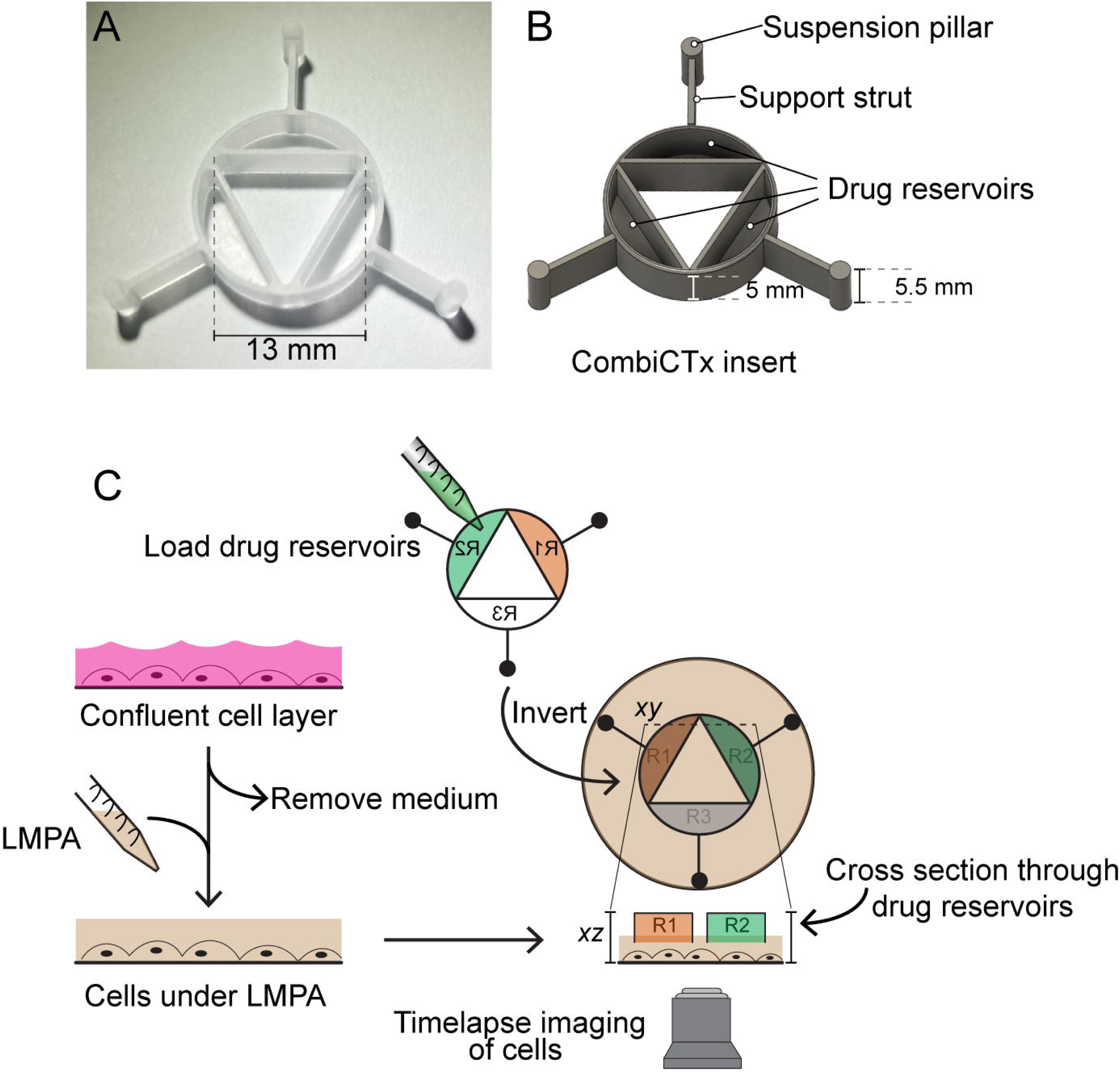
Overview of the CombiCTx insert and assay. **A.** Photograph of a CombiCTx insert. **B.** Computer-aided design rendering of the CombiCTx insert with relevant parts labelled. **C.** Overview of the CombiCTx assay. Cells are grown to confluence in a standard 6-well plate culture dish. Culture medium is aspirated and cells are covered in a layer of low-melting point agarose (LMPA). Reservoirs (R1-R3) in the CombiCTx insert are pre-filled with drugs of interest in polymerized LMPA. The CombiCTx insert is inverted and positioned over the LMPA-covered cells. The support struts help to centralize the insert within the well, and the suspension pillars ensure the insert is positioned a fixed distance above the cells. Cell responses are monitored by time-lapse confocal microscopy.

Depending on the specific conditions of the diffusion assays, the LMPA gel solution used to cover the cell layer contained combinations of the following cell stains: 2 µM propidium iodide (PI; Invitrogen Thermo Fisher Scientific), 2 µM Cell event caspase-3/7 green detection reagent (Thermo Fisher Scientific) and NucBlue live cell stain (Invitrogen Thermo Fisher Scientific), added in accordance with the manufacturer’s instructions.

### Fluorophore preparation for CombiCTx assays using FITC and TRITC

Fluorescein 5/6- isothiocyanate (FITC) or tetramethylrhodamine-5-isothiocyanate (TRITC) (Thermo Fisher Scientific, Uppsala, Sweden) were diluted to a concentration of 25 µM in 0.3% LMPA and added to separate adjacent reservoirs of a 3D-printed CombiCTx insert, and the remaining well was filled with 0.3% LMPA only.

### Drug preparation for CombiCTx assays using DOX, staurosporine, navitoclax and gemcitabine

For the drug diffusion assays, DOX (2 mg/ml; Accord Healthcare, Solna, Sweden; (Vnr: 189790); diluted in saline) was diluted to 100 µM, or staurosporine (Abcam, Cambridge, United Kingdom) was diluted to 50 µM in 0.6% LMPA and used to fill the desired number of reservoirs. For the combinatorial drug testing assays, navitoclax (CAS number 923564-51-6, Seleck Chemicals, Fisher Scientific) and gemcitabine (CAS number 122111-03-9, Seleck Chemicals, Fisher Scientific), were diluted to a concentration of 300 µM. To ensure consistent LMPA concentrations, a volume of DMSO equal to that used for the respective drugs was added to the LMPA used to fill the drug-free reservoirs.

### Physicochemical properties of the compounds used in the study

To facilitate discussions about potential differences in the diffusion of the studied molecules, the molecular mass; pKa; partition coefficient (Log P; a measure of a compound’s lipophilicity or hydrophobicity); and the distribution coefficient (Log D; which additionally accounts for the compound’s ionization state) are summarized in Table 1. The predicted Log P and Log D (at pH 7.4) values were collected from the Chemspider database (https://www.chemspider.com/) and were calculated using the PhysChem module of the ACD/Labs Percepta Platform software. Notably, predicted physicochemical values differ between databases depending on the calculation software used, and we provided the Chemspider records in Table 1 as it was the only source with Log P and Log D (at pH 7.4) values for all the molecules studied here, with the exception of TRITC. While we provide specific values in Table 1 we discuss differences in the lipophilicity between molecules and the potential effect on diffusion behavior in relative terms.

**Table 1.** Physicochemical properties of the compounds used in the study. Molecular weight (g/mol) were collected from PubChem (https://pubchem.ncbi.nlm.nih.gov/), as were the predicted pKa values for DOX, which cites the hazardous substances data bank as the data source. The remaining predicted pKa values were collected from the Drugbank database (https://go.drugbank.com/), which were calculated using Chemaxon software. The predicted Log P and Log D (at pH 7.4) values were collected from the Chemspider database (https://www.chemspider.com/), which were calculated with the PhysChem module of the ACD/Labs Percepta Platform software. All data was collected February 2025. (n/a; not available).

| Compound | Abbreviation | Molecular mass (g/mol) | pKa | Log P | Log D (at pH 7.4) |
| --- | --- | --- | --- | --- | --- |
| Tetramethylrhodamine-5-isothiocyanate | TRITC | 443.5 | n/a | n/a | n/a |
| Fluorescein 5/6-isothiocyanate | FITC | 389.4 | n/a | 4.00 | 4.64 |
| Doxorubicin | DOX | 543.5 | 7.34 (acidic, 1); 8.46 (2); 9.46 (basic, 3) | 2.82 | -0.79 |
| Staurosporine | STA | 466.5 | 13.46 (acidic), 9.55 (basic) | 3.29 | 3.30 |
| Gemcitabine | Gem | 263.2 | 11.52 (acidic), 3.65 (basic) | -0.47 | -1.40 |
| Navitoclax | Nav | 974.6 | 4.26 (acidic), 8.30 (basic) | 12.14 | 8.31 |

### Confocal microscopy, image acquisition and analysis

All imaging was performed on an LSM700 confocal laser scanning microscope (Zeiss, Jena, Germany) using a 5x/0,16 objective. Time-lapse images of fluorescently labelled cells, or fluorescent drugs were acquired using Zen imaging software (Zeiss, Jena, Germany). The tile scan function was applied to acquire 11 x 11 frame fields of view (approximately 23 mm x 23 mm), using a digital zoom setting of 0.6. The durations of the recordings are specified in the respective figures. Image analysis was performed using the Fiji version of ImageJ (22). For the doxorubicin and staurosporine data, the timelapse series were acquired in one continuous imaging session (using a temperature-controlled enclosure), whereas for the combinatorial drug experiments cells were returned to the incubator between the three timepoints (0, 24 and 48 hours), which were combined for each well. All images were then aligned in ImageJ using the rigid body transformation of the Hyperstack Registration plugin (23). Finally, 16 regions of interest (ROI) were positioned according to a predefined map, relative to specific insert features, to remain consistent across all repeats. Subsequently, the mean fluorescent intensity of each ROI in all three fluorescent channels was recorded for each well. For instances where the registration procedure failed to align the images satisfactorily, the 16 ROIs were positioned for each timepoint separately, and the analysis was performed as for all other wells.

We applied the Bliss independence model to calculate the expected combination effect (*E_Bliss_*) of navitoclax and gemcitabine based on the cell death results from the same ROIs obtained in assays using navitoclax only or gemcitabine only, and compared them to the results obtained in assays in which navitoclax and gemcitabine were combined (Fig. 6A-C). We applied the same approach to compare distinct ROIs within assays in which cells were treated with the combination of navitoclax and gemcitabine (Fig. 6D). We selected the Bliss independence model as it assumes that the drugs in question have distinct modes of action or targets (24), which is the case for navitoclax and gemcitabine, and was calculated using the formula:

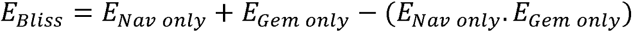

To compared the predicted combination effects (*E_Bliss_*) with the experimentally observed drug combination effects (*E_Nav+Gem_*) we calculated the combination index (CI):

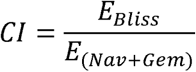

CI values that were below, equal to, or above 1 were respectively indicative of drug synergies, additive effects, or antagonisms (25).

### Statistical analyses

Statistical analyses were performed using Graphpad Prism 10 software. A Shapiro-Wilk test was used to test for normality of data distribution. For normally distributed data, an ordinary one-way ANOVA was applied to determine if there were significant differences between the means of groups (at P < 0.05). For the data from the DOX experiments a two-way repeated measures ANOVA was applied, and for both ANOVA types a Tukey’s multiple comparisons test identified significant differences between specific groups.

## Results and Discussion

The CombiCTx assay is based on the previously developed CombiANT assay, used for antibiotic interaction testing (26), and redesigned here for the purpose of studying the combined interactions and effects of anti-cancer drugs on adherent cancer cells. Central to the assay is the CombiCTx insert (Fig. 1A) that comprises three reservoirs, which can be filled with drugs of interest that are dissolved in low melting point agarose (LMPA). The CombiCTx insert features three support struts, each radiating outward from the midpoint of the outer wall of each reservoir, with a suspension pillar at their distal end that fixes a spatial distance of 0.5 mm from the bottom of the cell culture dish to the drug-loaded reservoir (Fig. 1B). The assay is activated in a two-step procedure: firstly, the CombiCTx reservoirs are loaded with the LMPA containing the drugs of interest and allowed to polymerize, next the cells are covered in a layer of LMPA, and the drug-loaded CombiCTx is inverted and gently lowered into the LMPA covering the cell layer prior to its polymerization (Fig. 1C). As the drug reservoirs face downwards towards the cells, once the LMPA covering the cells polymerizes it forms a continuous connection with the drug-containing LMPA in the reservoirs. This ensures that drugs can diffuse from each of the three drug reservoirs in a continuous gel towards the cancer cells, and thereby form dynamic gradient landscapes that expose cells to a wide range of drug concentrations for each of the studied drugs.

### Imaging patterns of combined fluorophore diffusion in the CombiCTx assay

Diffusion in hydrogels will be influenced by the molecular mass of the compounds of interest; potential interactions between the compounds and the diffusion matrix; and characteristics such as the pore size and molecular occupancy of the hydrogel (27–30). To visually assess and characterize the formation of dynamic gradients in the CombiCTx assay we imaged the diffusion of two fluorophores FITC and TRITC from two different reservoirs of the CombiCTx insert, outlined in Fig. 2A. FITC and TRITC have respective molecular mass of 389.4 g/mol and 443.5 g/mol, which are similar to commonly used anti-cancer drugs such as DOX (543.5 g/mol), and the reported pore sizes for agarose gels of a similar percentage to those used here are in the low micrometer range (31). Using Molview (https://molview.org/) we estimated the molecular diameters of FITC and TRITC to be respectively 1.25 nm and 1.46 nm, and so the relatively larger hydrogel mesh size is unlikely to have size-exclusion effects on fluorophore diffusion (27). The isothiocyanate group in both FITC and TRITC reacts with amine groups at pH 7–9, but as agarose gels are comprised of linear galactose-based polysaccharides with chemical formula (C_12_H_18_O_9_)_n_ (32), no such interactions are expected (33). Further, given that the average surface charge on agarose is predicted to be neutral (34), the diffusion of fluorophores would not be expected to be extensively electrostatically impeded. Together, this suggests that FITC and TRITC would diffuse efficiently in a similar manner in the LMPA gels. To visualize this a field of view incorporating the FITC- and TRITC-filled reservoirs was acquired by tile-scanning, which stitches together multiple acquisition frames (Fig. 2B). Signal reduction was observed at the edges between adjacent frames within the tile-scan, which is a recognized imaging artefact sometimes associated with this acquisition mode (35), so in an effort to smooth the transitions between frames we processed the tile-scanned images in ImageJ with a Gaussian blur filter (Fig. 2C). A montage of the time-lapse imaging for the fluorophores illustrates their diffusion patterns over a 14.5 h duration, over which they entered the central triangular interaction area (Fig. 2D). As the area below the apex of the triangle is closest to the FITC and TRITC reservoirs, it would be the first region in which diffusing FITC and TRITC will combine. To visualize this combination the fluorescence profile between the FITC and TRITC reservoirs was quantified using a line plot (white line in Fig. 2C) in the 0 h and 14.5 h images, which revealed the formation of overlapping and opposing FITC and TRITC gradients (Fig. 2E). To better visualize the temporal and spatial changes and overlap in FITC and TRITC diffusion, a threshold was set for the FITC and TRITC signals from the timelapse in Fig. 2D and the detectable moving front of these signals were outlined for each timepoint, and projected into a single FITC, TRITC and Merge image (Fig. 2F). From the earliest timepoint the combination of FITC and TRITC diffusion gradients between the adjacent FITC and TRITC reservoirs was apparent as an overlap of FITC and TRITC fronts in the upper region of the device. At later timepoints this overlap spread to the centre of the device. In regions where the FITC and TRITC reservoirs were adjacent to the lower empty reservoir no FITC and TRITC signal overlap was apparent, indicating that no combinatorial gradients were formed in these locations (Fig. 2F). To quantitatively assess these differences, the total FITC and TRITC fluorescence from three regions of interest (ROIs) were quantified. As expected, the combination of FITC and TRITC gradients in ROI(F+T), resulted in a higher degree of total fluorescence than in ROI(F) or ROI(T), in which only FITC or TRITC gradients had been observed (Fig. 2G). At the endpoint of the assay the total fluorescence observed in ROI(F+T) was similar, though slightly lower, than the sum of the fluorescence recorded for ROI(F) and ROI(T) (Fig. 2H). Together this data demonstrates the diffusion landscapes that can be formed between different compounds in the CombiCTx device, and highlights how distinct regions within the device can be used to compare combinatorial interactions and effects with those of individual compounds.

**Figure 2.**
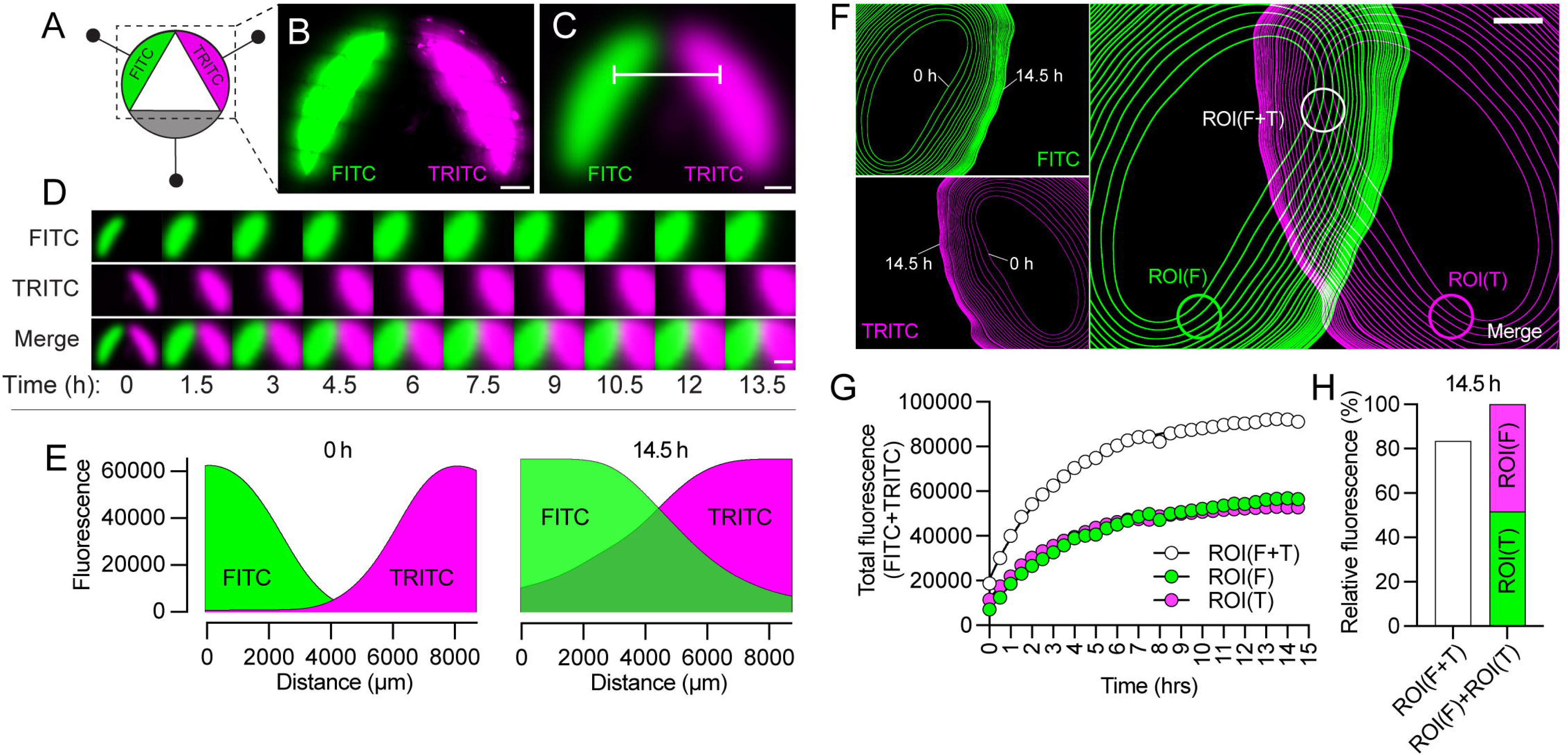
Timelapse imaging of FITC and TRITC diffusion gradients in the CombiCTx assay. **A.** Schematic illustration of the diffusion assay. The fluorophores FITC and TRITC were loaded in two adjacent reservoirs of the CombiCTx device. **B.** FITC and TRITC fluorescence signals at the 0 h timepoint from within the reservoirs of a CombiCTx device, acquired by tile-scanning. Scale bar; 2 mm. **C.** The image in B (and associated timelapse image stack) was subjected to the Gaussian blur processing function in ImageJ to smoothen the FITC and TRITC fluorescence signals acquired by tile scanning. **D.** Montage of the FITC and TRITC fluorescence from the CombiCTx insert in C, as imaged by timelapse confocal microscopy. Scale bar; 5 mm. **E.** Fluorescence profile plots of FITC and TRITC fluorescence at the start (0 h) and at the end (14.5 h) of the fluorophore diffusion assay. FITC and TRITC fluorescence were measured under the white line in C. **F.** Projections for the timelapse image stacks of the detectable diffusion fronts of FITC and TRITC fluorescence as they diffused from their respective CombiCTx reservoirs (see Material and Methods for details). For each fluorophore the innermost boundary line representing the 0 h timepoint, and the outermost boundary line representing the 14.5 h timepoint, is labelled. **G.** Quantification of the total fluorescence (FITC and TRITC) measured over time in the regions of interest (ROIs) identified in **F** as ROI(F+T), in which FITC and TRITC diffusion gradients were combined, and as ROI(F) and ROI(T), in which respectively FITC-only and TRITC-only fluorescence was observed. **H.** Comparison of the total fluorescence measured in ROI(F+T) compared to the sum of the total fluorescence measured in ROI(F) and ROI(T) at the 14. 5 h timepoint.

### Imaging doxorubicin (DOX) diffusion gradients in the CombiCTx assay

To characterize the formation of dynamic gradients of a small molecule anti-cancer drug within the CombiCTx assay we imaged the diffusion of DOX, a widely studied and clinically used (e.g. as a breast cancer treatment) anthracycline chemotherapeutic (36). DOX has a molecular mass of 543.5 g/mol, and its distinct photophysical properties permit quantification of drug concentrations by measuring absorbance or fluorescence (30, 37), which enabled us to visualize drug diffusion by tracking DOX fluorescence in the CombiCTx assay. The assay was conducted with LMPA-covered MDA-MB-231 cells, and the CombiCTx device was loaded without DOX (No DOX), with DOX loaded in reservoir 1, or DOX loaded in reservoirs 1 and 2 (Fig. 3A). DOX was imaged over 20 h by timelapse confocal microscopy and average fluorescence was quantified from 16 regions of interest, contained within the outer perimeter of the CombiCTx device (Fig. 3A). The positions were chosen to represent multiple areas directly under the three reservoirs (ROIs 4-5-6; 8-9-10 and 12-1-2), the interaction zones between adjacent reservoirs (ROIs 3-7-11), areas directly outside of the reservoir’s fronts (ROIs 13-14-15), and in the centre of the assay area (ROI-16). The DOX-free condition displayed a consistent level of minimal background fluorescence across all time points (Fig. 3A). In the conditions in which DOX was loaded in reservoir 1, or in reservoirs 1 and 2 simultaneously, the fluorescence in the reservoirs decreased over time as DOX diffused outwardly into the surrounding gel, and by the 20 h timepoint the fluorescence in the reservoirs approached the background signal as imaged in No DOX conditions (Fig. 3B). To quantitatively assess the cumulative DOX exposure, we calculated the area under the curve of DOX fluorescence for each ROI in each of the assays (Fig. 3C-E). To visualize the change in DOX levels as a result of diffusion, the signals from the centre of reservoir 1, through the centre of the device, and on to the centre of reservoir 2 were quantified, and the area under the curve (AUC) from the DOX signals collected over time in ROIs 5, 14, 16, 15 and 9 were plotted (Fig. 3C). As expected, when DOX was loaded only in reservoir 1, significant differences were detected between the selected ROIs that were positioned along the concentration gradient formed from the centre of the reservoir and inward towards the centre of the device. When DOX was loaded in reservoirs 1 and 2 symmetrical gradients of DOX were observed from each side of the device, and equal exposure was achieved in spatially equivalent positions (i.e. ROI-5 with ROI-9, and ROI-14 with ROI-15) (Fig. 3C). Next, we compared DOX exposure in ROIs that were placed within the perimeter of each of the reservoirs, and as close to an adjacent reservoir as possible (i.e. ROIs 2, 4, 6, 8, 10 and 12), to assess the degree to which diffusion between reservoirs contributes to DOX exposure within a given reservoir (Fig 3D). With DOX in reservoir 1, the accumulated fluorescence in the peripheral regions ROI-4 and ROI-6 were equal, as was the degree of DOX diffusion into the adjacent ROI-2 and ROI-8 in the DOX-free wells. When DOX was loaded in reservoir 1 and 2 the diffusion into ROI-2 and ROI-12 in the DOX-free reservoir was similar (Fig 3D). However, ROI-6 and ROI-8, which were both within and adjacent to a DOX reservoir, did not have significantly higher DOX levels than those recorded in ROI-4 or ROI-10, which were in contrast adjacent to DOX-free reservoirs (Fig. 3D). Finally, we investigated DOX exposure in the “interaction zones” between reservoirs *i.e.* ROI-3, ROI-7 and ROI-11 (Fig. 3E). As expected, when DOX was present only in reservoir 1, levels were highly similar between ROI-3 and ROI-7, and significantly lower in ROI-11 (Fig. 3E). Further, with DOX loaded both in reservoirs 1 and 2, the expected mass transfer of DOX would affect the levels for the interaction zone ROI-7 between DOX reservoirs, which was equal to the sum of the DOX levels quantified in ROI-3 and ROI-11, each of which were adjacent to the DOX-free reservoir (Fig. 3E). These data further support the observations in Fig. 2, which highlight the utility of analyzing interaction regions within the CombiCTx assay to quantify the combined effects of two drugs, while having internal control positions within the same device for assessing the interaction and response to individual drugs along dynamic concentration gradients, which bears similarities with how drugs diffuse in tissues. An *in vitro* assay that accounts for these diffusion effects could provide valuable translational knowledge for locoregional anti-cancer treatments such as transarterial chemoembolization and local prostate injections (38, 39).

**Figure 3.**
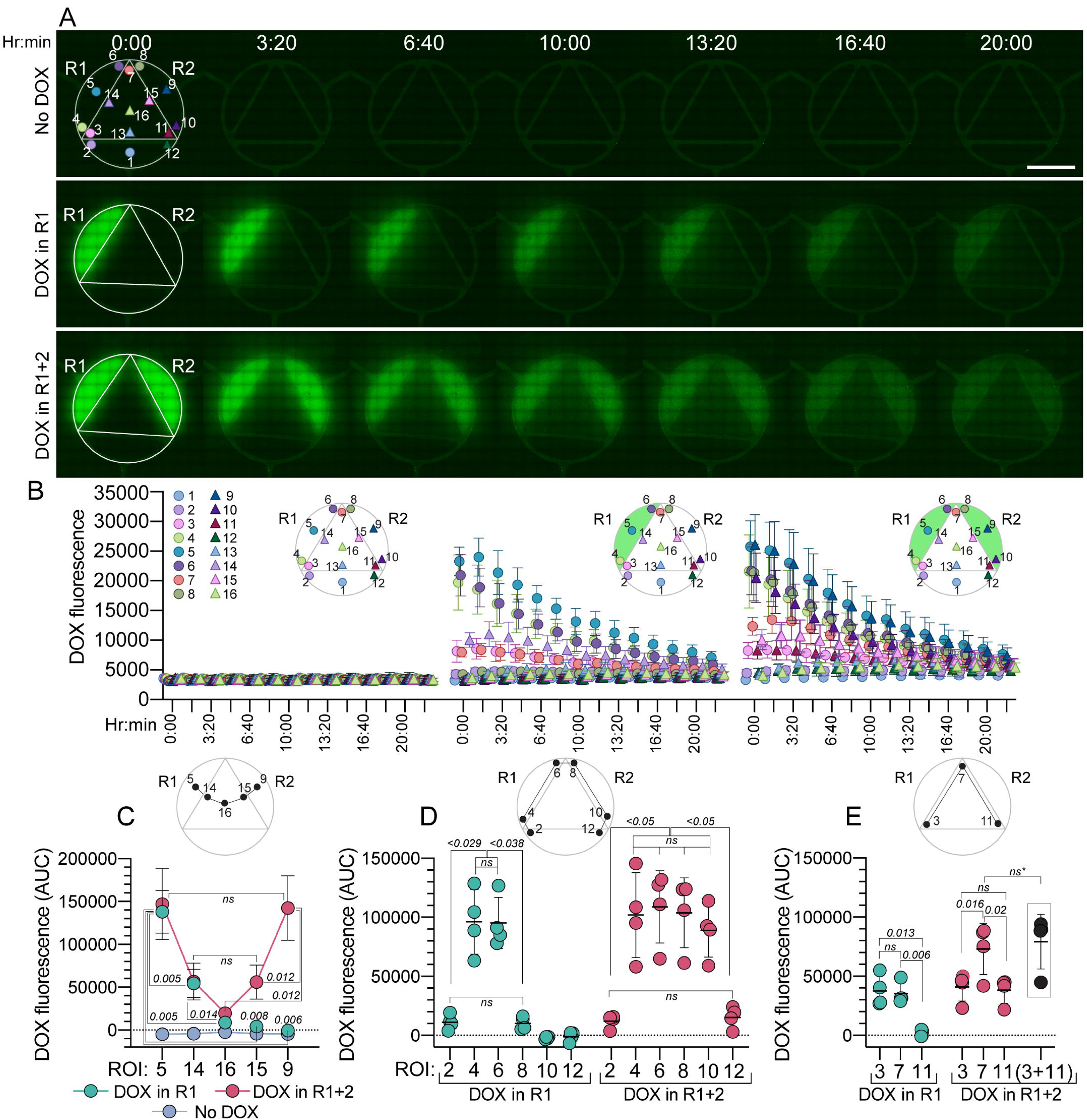
Doxorubicin diffusion in the CombiCTx assay. **A.** Tile-scanned overview of a CombiCTx assay in which no reservoirs were loaded with doxorubicin (DOX), DOX was loaded in reservoir 1 (R1), or DOX loaded in reservoirs 1 and 2 (R1+2) of the CombiCTx device and DOX fluorescence was imaged over time by fluorescence microscopy. Illustrations of the CombiCTx device are superimposed for orientation, to indicate reservoir 1 (R1) and reservoir 2 (R2), and the 16 regions of interest (ROIs), which have been quantified. Scale bar: 5 mm. **B.** Quantification of DOX fluorescence intensity for each of the conditions and timepoints presented in A, within each of the 16 ROIs identified above. Each point and error bars in the scatterplot represent the mean and standard deviation from four independent experiments. **C-E.** Quantification of the area under the curve (AUC) of DOX fluorescence presented in **B**, representing the accumulated DOX fluorescence for each of the indicated ROIs, for the duration of the experiment (20 h). The respective positions of the compared ROIs within the CombiCTx device are illustrated above each scatterplot. The points and error bars in the plots in **C** and the scatterplots in **D** and **E** represent the mean and standard deviation from four independent experiments. Significant differences between ROIs were evaluated by two-way repeated measures ANOVA, with a Tukey’s multiple comparisons test.

### Quantitative image analysis of apoptosis and cell death in the CombiCTx assay

To facilitate quantification of the response to anti-cancer agents using the CombiCTx assay, we next adapted an imaging protocol using a caspase 3/7 reporter to detect apoptotic cells and the nuclear stain propidium iodide (PI, which is impermeable to living cells) to detect dead cells. MDA-MB-231 breast cancer cells were pre-seeded on the bottom of a cell culture dish and cultured under standard conditions. Prior to initiating the CombiCTx assay, all three CombiCTx reservoirs were loaded with 50 µM staurosporine, which has a molecular mass of 466.5 g/mol, which is somewhat smaller than DOX (Table 1). Staurosporine is a potent protein kinase C inhibitor and an established inducer of apoptosis (40), capable of acting through both caspase-dependent and caspase-independent mechanisms (41). We have previously demonstrated that staurosporine induces detectable caspase 3/7 activation in MDA-MB-231 cells after approximately 5 h of exposure (42). The cancer cells were covered with an LMPA solution in which the caspase 3/7 reporter, PI, and the live nuclear stain NucBlue were diluted, and the staurosporine-loaded CombiCTx insert was then inverted and positioned in the dish to activate the assay. Time-lapse confocal microscopy imaging of labelled MDA-MB-231 cells under and around the entire CombiCTx insert was carried out to visualize the induction of apoptosis and occurrence of cell death over time (Fig 4A-C). Gradual caspase 3/7 activation and increased cell death (i.e. PI-positive cells) were primarily observed in the regions directly under or close to the drug reservoirs (Fig. 4A-C). After 10 h the majority of cells directly under the reservoir were positive for caspase 3/7 and PI, indicating widespread cell death (Fig. 4C area iii), while imaging at positions directly outside the same reservoir (Fig. 4C area ii), revealed more apoptotic cells, and further into the central most interaction space of the CombiCTx insert (Fig. 4C area i) revealed low levels of apoptosis and death, in line with the expected staurosporine concentration gradient. Image analysis of the average caspase 3/7 and PI signal intensities, relative to the NucBlue intensities was carried out in the previously defined 16 ROIs (Fig. 4D) to quantify the induction of apoptosis and cell death during the first 10 hours after assay activation (Fig. 4 E and F). Caspase 3/7 activity and PI signal increased over time, with the highest increase recorded in ROIs directly beneath the reservoirs (Fig. 4E and F, ROIs 1, 5 and 9), as expected for these regions exposed to the highest staurosporine concentrations.

**Figure 4.**
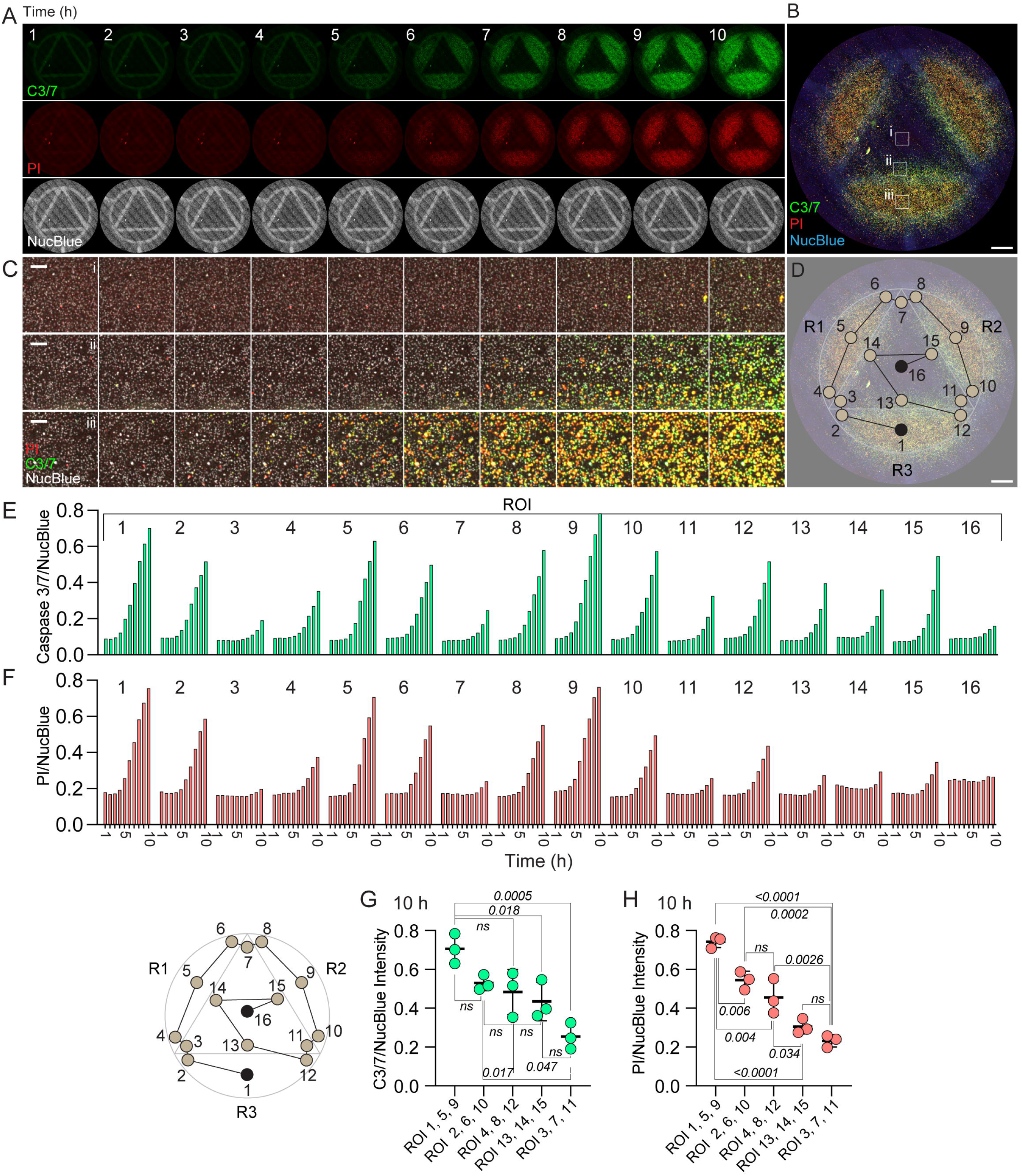
Assessment of apoptosis and cell death in distinct regions of interest in the CombiCTx assay. **A.** Tile-scanned overview of a 10 h CombiCTx assay in which MDA-MB-231 cells were exposed to staurosporine, which was loaded at a dose of 50 μM into each of the three CombiCTx reservoirs. MDA-MB-231 cell nuclei were labelled with NucBlue, apoptotic cells were identified using the Cellevent caspase 3/7 reporter, and dead cells were labelled by propidium iodide (PI) staining. Scale bar; 2 mm. **B.** Merged image of C3/7, PI and NucBlue fluorescence from the assay in A. at the 10 h timepoint. Scale bar; 2 mm. **C.** Enlarged views of the merged time-lapse images collected from the positions labelled in B as (i) an area in the center of the assay, (ii) an area close to the inner edge of the lower reservoir (R3), and (iii) an area from beneath R3. Scale bar; 250 μm. **D.** Regions of interest (ROIs) map overlaid on the image presented in **B**. Scale bar; 2 mm. **E.** Quantification of caspase 3/7 (C3/7) fluorescence normalized to the NucBlue fluorescence in each of the 16 ROIs for each timepoint. **F.** Quantification of propidium iodide (PI) fluorescence relative to NucBlue fluorescence in each of the 16 ROIs for each timepoint. **G and H.** Statistical comparison of ROIs grouped based on their positions relative to the centre of each reservoir for caspase 3/7 (C3/7)/NucBlue fluorescence (G), and propidium iodide (PI)/NucBlue fluorescence (H). ROIs 1, 5 and 9 are directly beneath the centre of each reservoir, while ROIs 2, 6 and 10 and ROIs 4, 8 and 12 are within, but furthest from the centre of their respective reservoirs. ROIs 13, 14, and 15 are directly outside the front of each reservoir, and ROIs 3, 7 and 11 represent regions between adjacent reservoirs. Datapoints, including the presented mean and standard deviations, represent values recorded at the 10 h timepoint from one CombiCTx device, and statistical comparisons were performed by one-way ANOVA with Tukey’s multiple comparisons test.

Quantitative comparisons between ROIs that were spatially matched relative to their proximity to the three staurosporine sources revealed similar caspase 3/7 activity (Fig. 4G) and PI staining (Fig. 4H) indicating that similar staurosporine diffusion gradients were established from each of the reservoirs. Therefore, in applications of the CombiCTx assay where the same drug is loaded in all reservoirs these matched ROIs can function as technical replicates. In general, caspase 3/7 activity and PI staining patterns were similar between respective ROIs; however, more significant differences were detected between the levels of PI staining than for caspase 3/7 activity. As expected, no differences were detected between the peripheral regions in opposing sides under each of the three reservoirs i.e. ROIs 2, 6 10 vs. ROIs 4, 8, 12, and though not statistically significant, relative caspase 3/7 activity and PI staining was lower in the interaction zone ROIs 3, 7, 11, than in ROIs 14, 15, 13, which were directly outside the midpoint of each reservoir (Fig. 4G and H). This was somewhat unexpected, as in Fig. 3 the additive levels of DOX measured in ROI-7 in experiments with DOX in both reservoirs (Fig. 3E) were higher than those recorded in ROIs 14 or 15 (Fig. 3C); therefore, we had expected a relatively higher degree of death in these zones where the staurosporine gradients from adjacent reservoirs converged. However, while staurosporine and DOX have relatively similar molecular mass other physiochemical properties may influence their respective diffusion in these regions. For example, the predicted Log D value (at pH 7.4) for staurosporine is higher than for DOX (Table 1), and this higher lipophilicity may reduce staurosporine diffusion in or impact its interactions with the agarose, but may also make it more cell permeable. Notably, an experimental determination of Log D (at pH 7.2) for DOX was higher (0.71) than the predicted value in Table 1 (−0.74 at pH 7.4), but still lower than staurosporine (43). Further, DOX has an established propensity to form self-associating aggregates (44), which would negatively impact diffusion, cell permeability and intracellular distribution. Nonetheless, this imaging strategy permits the quantitative assessment of anti-cancer drug effects on cell viability, and can be spatially and temporally resolved in the CombiCTx assay to study combinations of dynamic drug diffusion gradients.

### Assessing the cytotoxic effects of combinations of navitoclax and gemcitabine using the CombiCTx assay

Finally, we applied the CombiCTx assay to assess the cytotoxicity of combinations of two chemotherapeutic drugs with different physicochemical properties and mechanistic targets; namely, navitoclax and gemcitabine. Navitoclax is a Bcl2-inhibitor that has been investigated in phase I and II clinical trials (45). Gemcitabine, an approved chemotherapeutic treatment of the anti-metabolite class, inhibits DNA synthesis through integrating into the new DNA strand and inhibiting the nucleoside salvage pathway thus hindering DNA replication (46). Jaaks et al. reported robust synergy between a navitoclax and gemcitabine combination treatment of basal-like breast cancer cell lines (including MDA-MB-231 cells) in a 2D assay, with synergies detected in approximately 69% of the 23 breast cancer cell lines included in the validated subset (11). Navitoclax and gemcitabine have also been tested in a combinatorial dosing regimen in a phase I clinical trial in patients with solid tumors and aimed to determine safety, optimal dosing, and clinical activity. The regimen was well tolerated, but provided a best response of stable disease i.e. the extent and severity of the cancer neither increased or decreased (47).

The MDA-MB 231 cells were exposed to navitoclax-only, gemcitabine-only, or navitoclax + gemcitabine by loading one or two drug reservoirs in the same assay. Imaging was performed directly after activation of the CombiCTx assay and again after 24 h and 48 h, and was returned to the incubator between these timepoints. The apoptosis and cell death imaging protocol described in Fig. 4 was applied to determine the extent of cytotoxicity (Fig 5). An example of the collected tile scan images is presented in Fig. 5A. While the tile scan covers a large area with low magnification (5x objective), signals from single cells were readily detected (Fig. 5B). Quantification of relative caspase 3/7 activity and PI signals per NucBlue intensity was performed in the same 16 ROIs as described previously. Caspase 3/7 activity increased over time, as expected, with the highest activity recorded at 48 hrs in the navitoclax + gemcitabine assay in the region directly under the navitoclax drug reservoir (Fig. 5C). Similarly, the highest relative PI signal was recorded after 48 h under the navitoclax-filled drug reservoir (ROI-4, −5 and −6), in the navitoclax + gemcitabine assay (Fig. 5D). Overall, the variability in the relative caspase 3/7 intensity was high between replicates, while a greater degree of agreement was detected between ROI replicates for the relative PI staining (Fig. 5 C and D). Notably, at the 0 h timepoints we detected elevated PI fluorescence under wells containing navitoclax. As this timepoint is acquired directly after activation of the CombiCTx assay, the PI signal is not readily attributed to cell death, and to our knowledge navitoclax has no intrinsic fluorescence properties. Nonetheless, to ensure that subsequent quantitative analyses would only account for the degree to which the relative PI signal changed during the 48 h time period, we subtracted the 0 h timepoint intensities from the 48 h data for respective ROIs (Fig. 6).

**Figure 5.**
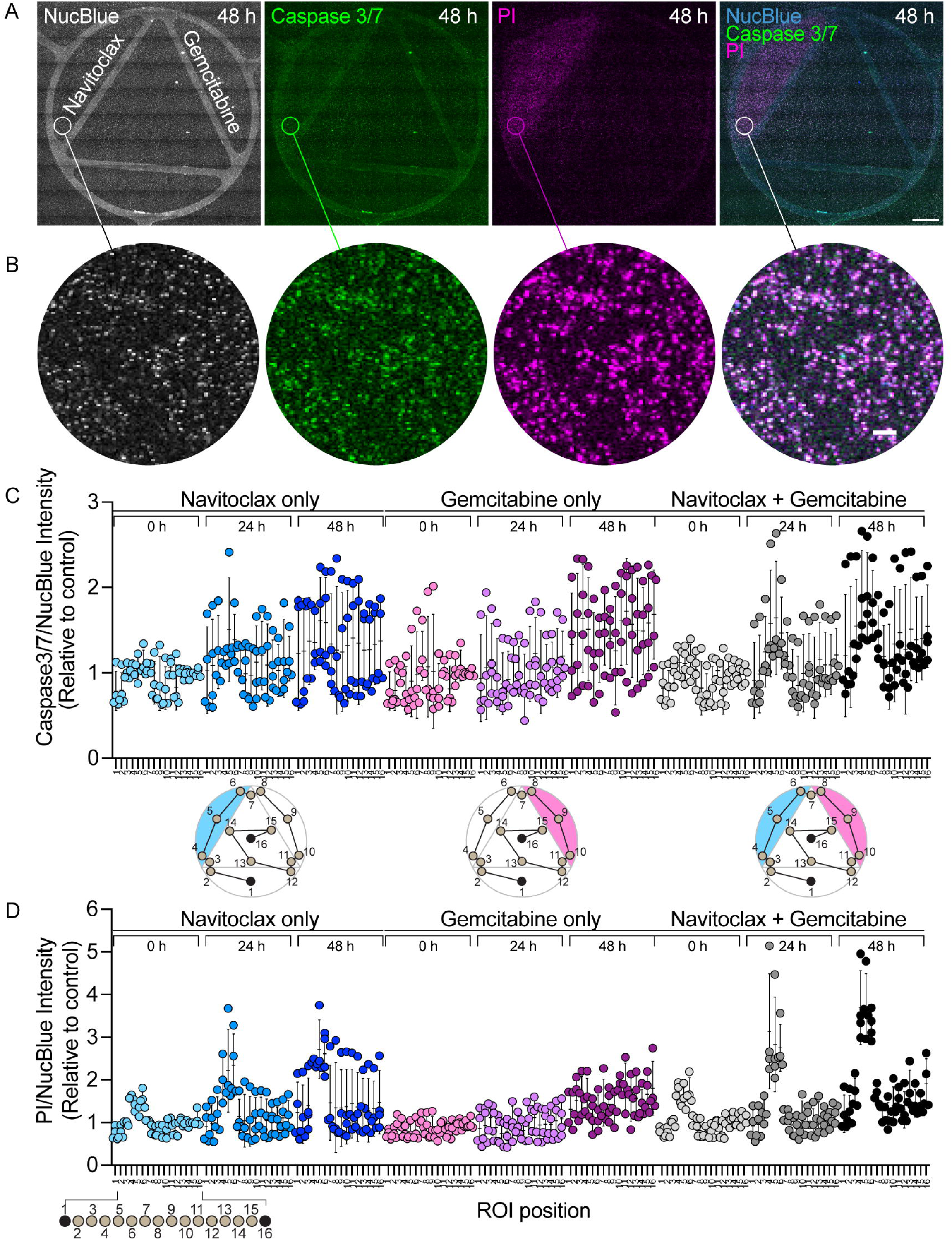
Assessment of navitoclax and gemcitabine cytotoxicity using the CombiCTx assay. **A.** Tile-scanned overview of a CombiCTx assay in which MDA-MB-231 cells were exposed to navitoclax and gemcitabine for 48 h and imaged by fluorescence microscopy. Cell nuclei were labelled with NucBlue, apoptotic cells were identified using the Cellevent caspase 3/7 reporter, and dead cells were labelled with propidium iodide (PI). Scale bar; 2 mm. **B.** Enlarged views of the respective areas encircled in A. Scale bar; 100 μm. **C.** Quantification of caspase3/7 fluorescence/NucBlue fluorescence intensity in each of the 16 regions of interest (relative positions indicated on the ROI maps below) for CombiCTx assays in which cells were treated with navitoclax only (loaded in the upper left reservoir), gemcitabine only (loaded in the upper right reservoir), or both navitoclax (left reservoir) and gemcitabine (right reservoir) for 0, 24 and 48 h. **D.** Quantification of PI fluorescence/NucBlue fluorescence intensity in each of the 16 regions of interest (relative positions indicated on the ROI maps above) for CombiCTx assays in which cells were treated with navitoclax only, gemcitabine only, or navitoclax and gemcitabine for 0, 24 and 48 h. All datapoints in **C** and **D** represent values from four independent experiments.

As the establishment of diffusion gradients requires time, locations within the CombiCTx assay that are closer to one drug source than the other can also be used to assess the effects of sequential treatments. We compared the relative PI signal intensity in ROI-5, centrally located in the reservoir in which navitoclax was loaded in the navitoclax-only, and navitoclax + gemcitabine conditions, and which was drug-free in the gemcitabine only conditions. In the navitoclax + gemcitabine assay, cells in this location would be first exposed to a high concentration of navitoclax, followed later by a relatively lower concentration of gemcitabine. The relative PI signal in ROI-5 was significantly higher in the navitoclax + gemcitabine assay compared to the gemcitabine-only assay, and while not significant it was also higher than that observed for navitoclax-only (Fig. 6A). This PI signal from the combined treatment assays were also marginally greater than the *E_Bliss ROI5_* values, which calculates the expected effect of combining these two drugs based on the effects recorded in the single drug assays (Fig. 6A, datapoints within the rectangle). Furthermore, the combination indices calculated from the E_Bliss ROI5_ values and the corresponding data for the effects (*E*) recorded in the navitoclax + gemcitabine assays (*E_Nav+Gem_*) were all below 1 (Fig. 6E), which was indicative of mildly synergistic drug effects. In contrast, comparisons between the centrally located ROI-16 for the different assays revealed no significant differences (Fig. 6B). If both drugs diffused at an equal rate from their respective reservoirs the combined drug concentration in ROI-16 would be predicted to be higher in the navitoclax + gemcitabine than in the navitoclax-only or gemcitabine-only assays. However, for the navitoclax-only conditions the relative PI signal was close to 0 for the majority of replicates, but slightly elevated for the gemcitabine-only condition and the navitoclax + gemcitabine condition (Fig. 6B). A similar pattern was observed in ROI-9, within the reservoir loaded with gemcitabine in the gemcitabine only and navitoclax + gemcitabine assay, and gemcitabine-free in the navitoclax-only assay (Fig. 6C).

**Figure 6.**
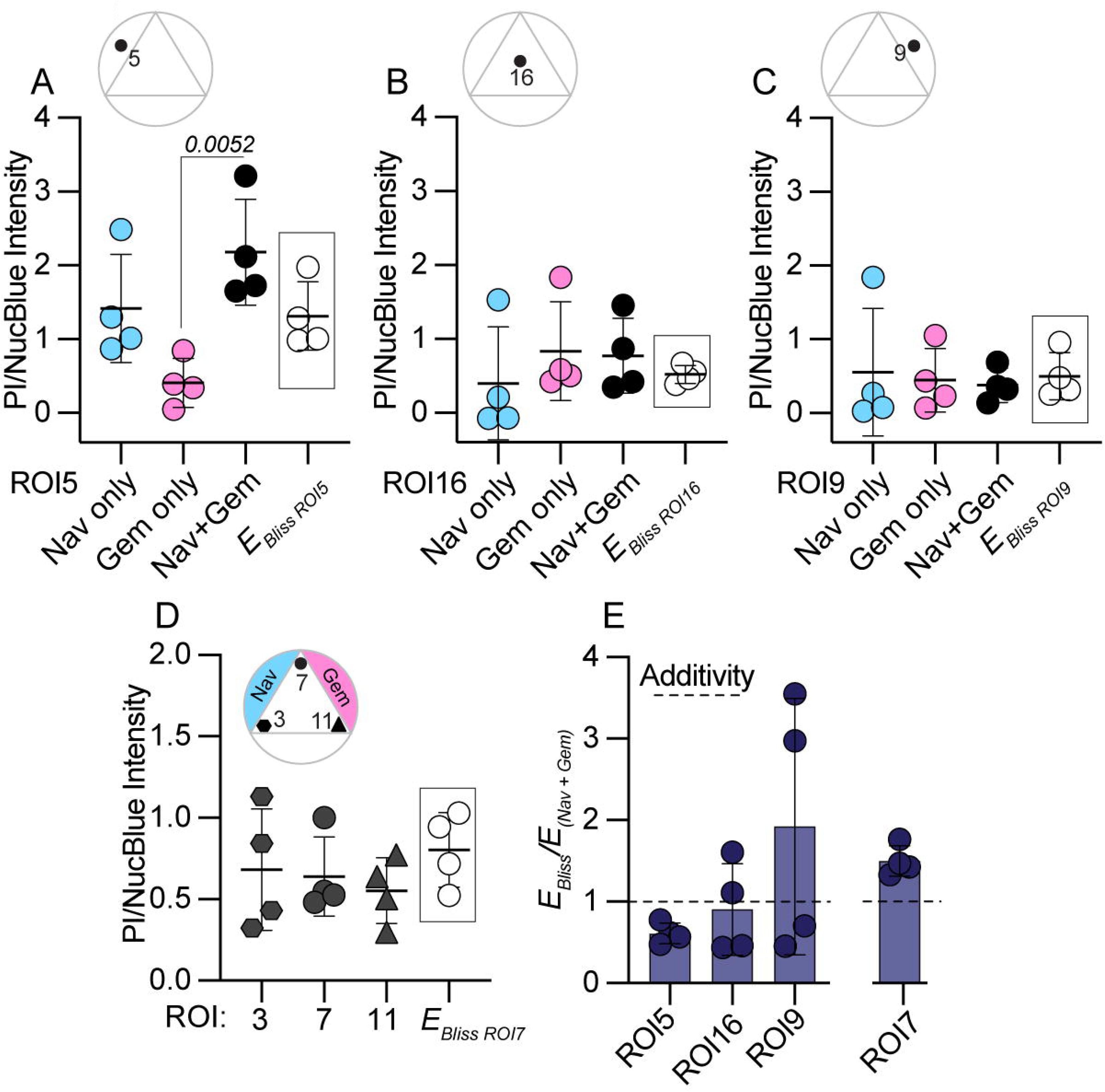
**A-C.** Inter-assay statistical comparisons of cell death as measured by propidium iodide (PI)/NucBlue intensity after 48 h (corrected for intensities at the 0 h timepoint) in specific ROIs (indicated on the respective ROI maps), from the assays outlined in Figure 5 where CombiCTx devices were loaded with navitoclax only (upper left reservoir), gemcitabine only (upper right reservoir) or navitoclax (Nav; left reservoir) and gemcitabine (Gem; right reservoir). For comparison with the Nav+Gem condition, the framed datapoints labelled as E_Bliss ROI5, ROI16, or ROI9_ represent the calculated drug combination effect, based on the Bliss independence model, for the PI/NucBlue intensities recorded in the Nav only and Gem only assays. **D.** Intra-assay statistical comparisons of cell death as measured by propidium iodide (PI)/NucBlue intensity after 48 h (corrected for intensities at the 0 h timepoint) in ROI 3, 7 and 11 (indicated on the associated ROI map), from the assays outlined in Figure 5 where CombiCTx devices were loaded with Nav and Gem. For comparison with ROI 7 (which is closest to both the Nav and Gem wells), the framed datapoints labelled as (E_Bliss ROI7_) represent the calculated drug combination effect, based on the Bliss independence model, for the PI/NucBlue intensities recorded in ROI 3 (which is closest to Nav and furthest from Gem) and ROI 11 (which is closest to Gem and furthest from Nav). **E.** Calculated indices (CI) of the predicted combination effects (*E_Bliss_*) compared to the experimentally observed drug combination effects (*E_Nav+Gem_*). Values that are below, equal to, or above 1 are respectively indicative of drug synergies, additive effects, or antagonisms. All datapoints, including the presented mean and standard deviations, represent values from four independent experiments. The significant difference in A was determined using a one-way ANOVA and a Tukey’s multiple comparisons test. No significant differences were detected between any of the other groups

This suggests that the degree of navitoclax diffusion within the assay is much lower than that of gemcitabine. The *E_Bliss_/E_Nav+Gem_* calculated for ROI16 and ROI9 were also highly variable, which may in part be due to the relatively low navitoclax concentrations reaching these locations. A possible explanation for these diffusion differences is the respective differences in the molecular mass and lipophilicity of these two drugs. Gemcitabine is the smallest molecule in our study (263.2 g/mol) (48), and is substantially less lipophilic than navitoclax (see Table 1) and has negligible protein binding affinities (49). In contrast, navitoclax was the largest drug tested in the assay with a molecular mass of 974.6 g/mol and high lipophilicity (see Table 1), with strong protein binding properties (50, 51). Therefore, it is likely that gemcitabine readily diffuses throughout the assay space impacting viability at a greater range, while navitoclax diffuses more slowly causing more localized effects, and may be further impeded by interactions with the cells seeded directly under the navitoclax reservoir. To compare the combined effect of navitoclax and gemcitabine exposure with the effect from predominantly navitoclax or gemcitabine alone within the same CombiCTx assay, we compared the relative PI signal recorded in ROI-3, ROI-7 and ROI-11. A similar degree of cell death was observed in all three locations, and the predicted combination effect (*E_Bliss ROI7_*) calculated by combining the cell death observed in ROI-3 (nearest navitoclax) with that in ROI-11 (nearest gemcitabine) was not observed in ROI-7 (which is equally near the navitoclax and gemcitabine reservoirs; Fig. 6D). As a result, the *E_Bliss_/E_Nav+Gem_*indices in ROI7 were above 1, indicating a mildly antagonistic effect of the combined drugs in this location (Fig. 6E).

Together these observations highlight the importance of complimenting standard 2D drug tests with 3D assays in which the effect of the extracellular environment on drug diffusion, other potential binding interactions, and relative differences in the sequence of drug exposures may be investigated. The CombiCTx assay provides one platform for conducting such studies. Notably, the manner in which cells are exposed to drugs in the CombiCTx assay represents some important similarities with how drugs reach cancer cells in the body, namely by diffusion to form dynamic and transient gradients through both normal and tumor tissue. Therefore, it would be interesting to apply the CombiCTx assay to evaluate how gradients are formed in different types of tissue-like matrices known to modulate drug responses (30, 52–56), and to determine how cells respond to and interact with drugs in these specific environments. Furthermore, adapting a high content imaging protocol such as Cell Painting to phenotypically profile cells within gradients of drugs and other effector combinations may increase the detection rate and types of cellular responses that could be identified within the CombiCTx assay (57–59).

## Acknowledgements

This study was conducted as part of the Additive Manufacturing for the Life Sciences (AM4Life) consortium, which is funded by Sweden’s Innovation Agency VINNOVA (grant number 2019-00029), and by additional grants to J. K. from the Swedish Cancer Foundation (Cancerfonden; 20 1285 PjF and 23 2692 Pj 01 H), and the Göran Gustafsson Foundation (Göran Gustafssons Stiftelser). 3D printing was performed at U-PRINT: Uppsala University’s 3D-printing facility at the Disciplinary Domain of Medicine and Pharmacy and SciLifeLab Uppsala. We would like to thank Johan Öhman from the Dept. of Cell and Molecular Biology, Uppsala University for his assistance with the CNC milling of the CombiCTx inserts. F. H. is supported by the Swedish Cancer Foundation (23 2776 Pj), the Swedish Society for Medical Research (grant number S17-0092), The Swedish Research Council (grant number 2021-01628) and Göran Gustafsson Foundation (Göran Gustafssons Stiftelser). H. L. is supported by the Swedish Cancer Foundation (Cancerfonden, grant number CAN 24 3519 Pj 01 H) and Swedish Research Council (grant numbers 2020-02367 and 2024-03166).

## Author contributions

Conceptualization: C.S., O.E., J.K., P.O’C. Methodology: C.S., A.L.C., S.B., B.S., O.E., O.D., H.L., F.H., J.K., P.O’C. Validation: C.S., A.L.C., S.C., B.S., O.E., O.D. Investigation: C.S., A.L.C., S.B., B.S., O.E., O.D., P.O’C. Formal analysis: C.S., A.L.C., P.O’C. Resources: H.L., F.H., J.K. Data curation: C.S., A.L.C., P.O’C. Visualization: C.S., A.L.C., J.K., P.O’C. Writing original draft: C.S., J.K., P.O’C. Writing – review and editing: C.S., A.L.C., S.C., B.S., O.E., O.D., H.L., F.H., J.K., P.O’C. Supervision: H.L., F.H., J.K, P.O’C. Funding acquisition: H.L., F.H., J.K.

## References

1. Hanahan D. Hallmarks of Cancer: New Dimensions. Cancer Discov. 2022 Jan;12(1):31–46. PubMed PMID: 35022204.

2. Hanahan D, Weinberg RA. Hallmarks of cancer: the next generation. Cell. 2011 Mar 4;144(5):646–74. PubMed PMID: 21376230.

3. Negrini S, Gorgoulis VG, Halazonetis TD. Genomic instability--an evolving hallmark of cancer. Nat Rev Mol Cell Biol. 2010 Mar;11(3):220–8. PubMed PMID: 20177397.

4. Lopez JS, Banerji U. Combine and conquer: challenges for targeted therapy combinations in early phase trials. Nat Rev Clin Oncol. 2017 Jan;14(1):57–66. PubMed PMID: 27377132. PMCID: PMC6135233. Epub 20160705.

5. Menden MP, Wang D, Mason MJ, Szalai B, Bulusu KC, Guan Y, et al. Community assessment to advance computational prediction of cancer drug combinations in a pharmacogenomic screen. Nat Commun. 2019 Jun 17;10(1):2674. PubMed PMID: 31209238. PMCID: PMC6572829. Epub 20190617.

6. Hwangbo H, Chae S, Kim W, Jo S, Kim GH. Tumor-on-a-chip models combined with mini-tissues or organoids for engineering tumor tissues. Theranostics. 2024;14(1):33–55. PubMed PMID: 38164155. PMCID: PMC10750204. Epub 20240101. eng.

7. Ianevski A, Giri AK, Gautam P, Kononov A, Potdar S, Saarela J, et al. Prediction of drug combination effects with a minimal set of experiments. Nat Mach Intell. 2019 Dec;1(12):568–77. PubMed PMID: 32368721. PMCID: PMC7198051. Epub 20191209.

8. Narayan RS, Molenaar P, Teng J, Cornelissen FMG, Roelofs I, Menezes R, et al. A cancer drug atlas enables synergistic targeting of independent drug vulnerabilities. Nat Commun. 2020 Jun 10;11(1):2935. PubMed PMID: 32523045. PMCID: PMC7287046. Epub 20200610.

9. Hwangbo H, Patterson SC, Dai A, Plana D, Palmer AC. Additivity predicts the efficacy of most approved combination therapies for advanced cancer. Nature Cancer. 2023 2023/12/01;4(12):1693–704.

10. Peters K, Lerma Clavero A, Kullenberg F, Kopsida M, Dahlgren D, Heindryckx F, et al. Melatonin mitigates chemotherapy-induced small intestinal atrophy in rats and reduces cytotoxicity in murine intestinal organoids. PLoS One. 2024;19(9):e0307414. PubMed PMID: 39226257. PMCID: PMC11371236. Epub 20240903. eng.

11. Jaaks P, Coker EA, Vis DJ, Edwards O, Carpenter EF, Leto SM, et al. Effective drug combinations in breast, colon and pancreatic cancer cells. Nature. 2022 Mar;603(7899):166-73. PubMed PMID: 35197630. PMCID: PMC8891012. Epub 20220223.

12. Binnewies M, Roberts EW, Kersten K, Chan V, Fearon DF, Merad M, et al. Understanding the tumor immune microenvironment (TIME) for effective therapy. Nat Med. 2018 May;24(5):541–50. PubMed PMID: 29686425. PMCID: PMC5998822. Epub 20180423.

13. Kreuger J, Phillipson M. Targeting vascular and leukocyte communication in angiogenesis, inflammation and fibrosis. Nat Rev Drug Discov. 2016 Feb;15(2):125–42. PubMed PMID: 26612664. Epub 20151127.

14. Rodrigues J, Heinrich MA, Teixeira LM, Prakash J. 3D In Vitro Model (R)evolution: Unveiling Tumor-Stroma Interactions. Trends Cancer. 2021 Mar;7(3):249–64. PubMed PMID: 33218948. Epub 20201118.

15. Datta P, Dey M, Ataie Z, Unutmaz D, Ozbolat IT. 3D bioprinting for reconstituting the cancer microenvironment. NPJ Precis Oncol. 2020;4:18. PubMed PMID: 32793806. PMCID: PMC7385083. Epub 20200727.

16. Wang Y, Jeon H. 3D cell cultures toward quantitative high-throughput drug screening. Trends Pharmacol Sci. 2022 Jul;43(7):569–81. PubMed PMID: 35504760. Epub 20220430.

17. Choo N, Ramm S, Luu J, Winter JM, Selth LA, Dwyer AR, et al. High-Throughput Imaging Assay for Drug Screening of 3D Prostate Cancer Organoids. SLAS Discov. 2021 Oct;26(9):1107–24. PubMed PMID: 34111999. PMCID: PMC8458687. Epub 20210611.

18. Brancato V, Oliveira JM, Correlo VM, Reis RL, Kundu SC. Could 3D models of cancer enhance drug screening? Biomaterials. 2020 Feb;232:119744. PubMed PMID: 31918229. Epub 20191226.

19. Fevre R, Mary G, Vertti-Quintero N, Durand A, Tomasi RF, Del Nery E, et al. Combinatorial drug screening on 3D Ewing sarcoma spheroids using droplet-based microfluidics. iScience. 2023 May 19;26(5):106651. PubMed PMID: 37168549. PMCID: PMC10165258. Epub 20230412. eng.

20. Chou TC. Preclinical versus clinical drug combination studies. Leuk Lymphoma. 2008 Nov;49(11):2059–80. PubMed PMID: 19021049. eng.

21. Yu M, Selvaraj SK, Liang-Chu MM, Aghajani S, Busse M, Yuan J, et al. A resource for cell line authentication, annotation and quality control. Nature. 2015 Apr 16;520(7547):307–11. PubMed PMID: 25877200.

22. Schindelin J, Arganda-Carreras I, Frise E, Kaynig V, Longair M, Pietzsch T, et al. Fiji: an open-source platform for biological-image analysis. Nature Methods. 2012 2012/07/01;9(7):676–82.

23. Sharma V. ImageJ plugin hyperstackreg V5. 6. Zenodo https://zenodo.org/record/2252521#YzagunZBz-g. 2018.

24. Duarte D, Vale N. Evaluation of synergism in drug combinations and reference models for future orientations in oncology. Current Research in Pharmacology and Drug Discovery. 2022 2022/01/01/;3:100110.

25. Foucquier J, Guedj M. Analysis of drug combinations: current methodological landscape. Pharmacology Research & Perspectives. 2015;3(3):e00149.

26. Fatsis-Kavalopoulos N, Roemhild R, Tang P-C, Kreuger J, Andersson DI. CombiANT: Antibiotic interaction testing made easy. PLOS Biology. 2020;18(9):e3000856.

27. Lieleg O, Ribbeck K. Biological hydrogels as selective diffusion barriers. Trends in Cell Biology. 2011 2011/09/01/;21(9):543-51.

28. Lieleg O, Baumgärtel RM, Bausch AR. Selective Filtering of Particles by the Extracellular Matrix: An Electrostatic Bandpass. Biophysical Journal. 2009 2009/09/16/;97(6):1569–77.

29. Pluen A, Netti PA, Jain RK, Berk DA. Diffusion of macromolecules in agarose gels: comparison of linear and globular configurations. Biophys J. 1999 Jul;77(1):542–52. PubMed PMID: 10388779. PMCID: PMC1300351. eng.

30. Degerstedt O, O’Callaghan P, Clavero AL, Gråsjö J, Eriksson O, Sjögren E, et al. Quantitative imaging of doxorubicin diffusion and cellular uptake in biomimetic gels with human liver tumor cells. Drug Deliv Transl Res. 2024 Apr;14(4):970–83. PubMed PMID: 37824040. PMCID: PMC10927899. Epub 20231012. eng.

31. Moore MJ, Bodera F, Hernandez C, Shirazi N, Abenojar E, Exner AA, et al. The dance of the nanobubbles: detecting acoustic backscatter from sub-micron bubbles using ultra-high frequency acoustic microscopy. Nanoscale. 2020;12(41):21420–8.

32. Nosrati H, Khodaei M, Alizadeh Z, Banitalebi-Dehkordi M. Cationic, anionic and neutral polysaccharides for skin tissue engineering and wound healing applications. International Journal of Biological Macromolecules. 2021 2021/12/01/;192:298–322.

33. Hermanson GT. Chapter 10 - Fluorescent Probes. In: Hermanson GT, editor. Bioconjugate Techniques (Third Edition). Boston: Academic Press; 2013. p. 395–463.

34. Kadlec M, Pekař M, Smilek J. Mechanical properties of agarose hydrogels tuned by amphiphilic structures. Colloids and Surfaces A: Physicochemical and Engineering Aspects. 2024 2024/11/05/;700:134791.

35. Legesse FB, Chernavskaia O, Heuke S, Bocklitz T, Meyer T, Popp J, et al. Seamless stitching of tile scan microscope images. Journal of Microscopy. 2015;258(3):223–32.

36. Bisht A, Avinash D, Sahu KK, Patel P, Das Gupta G, Kurmi BD. A comprehensive review on doxorubicin: mechanisms, toxicity, clinical trials, combination therapies and nanoformulations in breast cancer. Drug Deliv Transl Res. 2024 Jun 17. PubMed PMID: 38884850. Epub 20240617. eng.

37. Degerstedt O, Gråsjö J, Norberg A, Sjögren E, Hansson P, Lennernäs H. Drug diffusion in biomimetic hydrogels: importance for drug transport and delivery in non-vascular tumor tissue. European Journal of Pharmaceutical Sciences. 2022 2022/05/01/;172:106150.

38. Lilienberg E, Dubbelboer IR, Karalli A, Axelsson R, Brismar TB, Ebeling Barbier C, et al. In Vivo Drug Delivery Performance of Lipiodol-Based Emulsion or Drug-Eluting Beads in Patients with Hepatocellular Carcinoma. Molecular Pharmaceutics. 2017 2017/02/06;14(2):448–58.

39. Sjögren E, Tammela TL, Lennernäs B, Taari K, Isotalo T, Malmsten L-Å, et al. Pharmacokinetics of an Injectable Modified-Release 2-Hydroxyflutamide Formulation in the Human Prostate Gland Using a Semiphysiologically Based Biopharmaceutical Model. Molecular Pharmaceutics. 2014 2014/09/02;11(9):3097–111.

40. Bertrand R, Solary E, O’Connor P, Kohn KW, Pommier Y. Induction of a Common Pathway of Apoptosis by Staurosporine. Experimental Cell Research. 1994 1994/04/01/;211(2):314–21.

41. Belmokhtar CA, Hillion J, Ségal-Bendirdjian E. Staurosporine induces apoptosis through both caspase-dependent and caspase-independent mechanisms. Oncogene. 2001 2001/06/01;20(26):3354–62.

42. O’Callaghan P, Engberg A, Eriksson O, Fatsis-Kavalopoulos N, Stelzl C, Sanchez G, et al. Piezo1 activation attenuates thrombin-induced blebbing in breast cancer cells. Journal of Cell Science. 2022;135(7).

43. Formariz TP, Sarmento VHV, Silva-Junior AA, Scarpa MV, Santilli CV, Oliveira AG. Doxorubicin biocompatible O/W microemulsion stabilized by mixed surfactant containing soya phosphatidylcholine. Colloids and Surfaces B: Biointerfaces. 2006 2006/08/01/;51(1):54–61.

44. Menozzi M, Valentini L, Vannini E, Arcamone F. Self-Association of Doxorubicin and Related Compounds in Aqueous Solution. Journal of Pharmaceutical Sciences. 1984 1984/06/01/;73(6):766–70.

45. Mohamad Anuar NN, Nor Hisam NS, Liew SL, Ugusman A. Clinical Review: Navitoclax as a Pro-Apoptotic and Anti-Fibrotic Agent. Front Pharmacol. 2020;11:564108. PubMed PMID: 33381025. PMCID: PMC7768911. Epub 20201126. eng.

46. Alvarellos ML, Lamba J, Sangkuhl K, Thorn CF, Wang L, Klein DJ, et al. PharmGKB summary: gemcitabine pathway. Pharmacogenet Genomics. 2014 Nov;24(11):564–74. PubMed PMID: 25162786. PMCID: PMC4189987. eng.

47. Cleary JM, Lima CMSR, Hurwitz HI, Montero AJ, Franklin C, Yang J, et al. A phase I clinical trial of navitoclax, a targeted high-affinity Bcl-2 family inhibitor, in combination with gemcitabine in patients with solid tumors. Investigational New Drugs. 2014 2014/10/01;32(5):937-45.

48. 48. https://www.ebi.ac.uk/chembl/compound_report_card/CHEMBL888/. [Available from: https://www.ebi.ac.uk/chembl/compound_report_card/CHEMBL888/.

49. Liston DR, Davis M. Clinically Relevant Concentrations of Anticancer Drugs: A Guide for Nonclinical Studies. Clin Cancer Res. 2017 Jul 15;23(14):3489–98. PubMed PMID: 28364015. PMCID: PMC5511563. Epub 20170331. eng.

50. Choo EF, Boggs J, Zhu C, Lubach JW, Catron ND, Jenkins G, et al. The role of lymphatic transport on the systemic bioavailability of the Bcl-2 protein family inhibitors navitoclax (ABT-263) and ABT-199. Drug Metab Dispos. 2014 Feb;42(2):207–12. PubMed PMID: 24212376. Epub 20131108. eng.

51. Wendt MD. The Discovery of Navitoclax, a Bcl-2 Family Inhibitor. In: Wendt MD, editor. Protein-Protein Interactions. Berlin, Heidelberg: Springer Berlin Heidelberg; 2012. p. 231–58.

52. Asmani M, Velumani S, Li Y, Wawrzyniak N, Hsia I, Chen Z, et al. Fibrotic microtissue array to predict anti-fibrosis drug efficacy. Nature Communications. 2018 2018/05/25;9(1):2066.

53. Fernández-Colino A, Iop L, Ventura Ferreira MS, Mela P. Fibrosis in tissue engineering and regenerative medicine: treat or trigger? Advanced Drug Delivery Reviews. 2019 2019/06/01/;146:17–36.

54. Calitz C, Rosenquist J, Degerstedt O, Khaled J, Kopsida M, Fryknäs M, et al. Influence of extracellular matrix composition on tumour cell behaviour in a biomimetic in vitro model for hepatocellular carcinoma. Sci Rep. 2023 Jan 13;13(1):748. PubMed PMID: 36639512. PMCID: PMC9839216. Epub 20230113. eng.

55. Hussey GS, Dziki JL, Badylak SF. Extracellular matrix-based materials for regenerative medicine. Nature Reviews Materials. 2018 2018/07/01;3(7):159–73.

56. Pickup MW, Mouw JK, Weaver VM. The extracellular matrix modulates the hallmarks of cancer. EMBO Rep. 2014 Dec;15(12):1243–53. PubMed PMID: 25381661. PMCID: PMC4264927. Epub 20141108. eng.

57. Bray M-A, Singh S, Han H, Davis CT, Borgeson B, Hartland C, et al. Cell Painting, a high-content image-based assay for morphological profiling using multiplexed fluorescent dyes. Nature Protocols. 2016 2016/09/01;11(9):1757–74.

58. Rietdijk J, Aggarwal T, Georgieva P, Lapins M, Carreras-Puigvert J, Spjuth O. Morphological profiling of environmental chemicals enables efficient and untargeted exploration of combination effects. Sci Total Environ. 2022 Aug 1;832:155058. PubMed PMID: 35390365. Epub 20220404. eng.

59. Seal S, Trapotsi M-A, Spjuth O, Singh S, Carreras-Puigvert J, Greene N, et al. A Decade in a Systematic Review: The Evolution and Impact of Cell Painting. bioRxiv. 2024:2024.05.04.592531.

